# A Neurogenic Signature Involving Monoamine Oxidase-A controls Human Thermogenic Adipose Tissue Development

**DOI:** 10.1101/2021.12.29.474474

**Authors:** Javier Solivan-Rivera, Zinger Yang Loureiro, Tiffany DeSouza, Anand Desai, Qin Yang, Raziel Rojas-Rodriguez, Pantos Skritakis, Shannon Joyce, Denise Zhong, Tammy Nguyen, Silvia Corvera

**Author notes:** Corresponding author at, Program in Molecular Medicine, 373 Plantation Street, Worcester, MA 01605.

## Abstract

Mechanisms that control “beige/brite” thermogenic adipose tissue development may be harnessed to improve human metabolic health. To define these mechanisms, we developed a species-hybrid model in which human mesenchymal progenitor cells were used to develop white or thermogenic/beige adipose tissue in mice. The hybrid adipose tissue developed distinctive features of human adipose tissue, such as larger adipocyte size, despite its neurovascular architecture being entirely of murine origin. Thermogenic adipose tissue recruited a denser, qualitatively distinct vascular network, differing in genes mapping to circadian rhythm pathways, and denser sympathetic innervation. The enhanced thermogenic neurovascular network was associated with human adipocyte expression of THBS*4*, *TNC*, *NTRK3* and *SPARCL1*, which enhance neurogenesis, and decreased expression of *MAOA* and *ACHE*, which control neurotransmitter tone. Systemic inhibition of MAOA, which is present in human but absent in mouse adipocytes, induced browning of human but not mouse adipose tissue, revealing the physiological relevance of this pathway. Our results reveal species-specific cell type dependencies controlling the development of thermogenic adipose tissue and point to human adipocyte MAOA as a potential target for metabolic disease therapy.

## INTRODUCTION

A positive association between thermogenic “beige/brite” adipose tissue and metabolic health in humans has been repeatedly observed [1–4], and multiple studies have shown that human thermogenic adipocytes transplanted into mice affect whole-body metabolism [5–7]. These studies provide evidence for a cause-effect relationship and a rationale for enhancing the activity and/or abundance of thermogenic adipose tissue as a therapeutic approach to metabolic disease. However, the cellular and molecular mechanisms by which human thermogenic adipose tissue develops and is maintained are largely unknown.

In adults, thermogenic adipose tissue develops in response to chronic cold exposure, extensive skin burns, and catecholamine producing tumors [8–12]. Formation of new thermogenic adipose tissue involves the generation of new adipocytes expressing the characteristic mitochondrial uncoupling protein 1 (UCP1) of new vasculature to facilitate oxygen consumption and heat dissipation [13–16], and of new innervation to mediate sympathetic signaling [13]. Signaling between multiple cell types is likely to be involved in the process of tissue expansion; for example, adipocytes secrete factors that induce vascularization and innervation [17], catecholamines can stimulate angiogenesis, and progenitor cells niched in the newly forming vasculature can differentiate into thermogenic adipocytes [18–20]. The concurrence of adipogenesis, angiogenesis and innervation make it difficult to determine cell-type specific mechanisms and temporal interdependencies, and to identify those mechanisms that could be harnessed to enhance thermogenic adipose tissue mass or activity. Understanding mechanisms of human thermogenic adipose tissue expansion is particularly difficult due to limited models available to study human tissue development. Our laboratory has developed methods to obtain large numbers of mesenchymal progenitor cells from human adipose tissue and shown that these can give rise to multiple adipocyte subtypes [19]. Moreover, these cells can be implanted into immunocompromised mice, where they develop metabolically integrated adipose tissue [21]. We reasoned that this cell implantation model, by clearly separating the adipocyte component(human) from vascular and neuronal components (mouse), could help deconvolve mechanisms of development and maintenance of human thermogenic adipose tissue.

Here we report that human adipose tissue generated in mice maintains critical species-specific features, including human adipocyte cell size, thermogenic features, and the capacity to recruit a functional neurovascular network. Thermogenic adipocytes recruit a denser neurovascular network with distinct transcriptomic features, demonstrating that thermogenic adipose tissue vasculature is both quantitative and qualitatively different. The recruitment of a distinct neuro-vasculature by thermogenic adipocytes is directly associated with expression of genes involved in paracrine signaling of neurogenesis, and in modulation of neurotransmitter tone. Notably, *THBS4* and *SPARCL1*, which directly stimulate synapse formation [22], are the most highly differentially expressed genes distinguishing thermogenic from non-thermogenic adipocytes in the hybrid tissue. In addition, expression of MAOA, which catalyzes the oxidative deamination and inactivation of multiple neurotransmitters [23], is suppressed in adipocytes that give rise to thermogenic tissue. Functionally, systemic inhibition of MAOA with the irreversible inhibitor clorgyline results in beiging of hybrid adipose tissue, but not of endogenous mouse adipose tissue. These data reveal species-distinct mechanisms by which adipocytes establish the specialized neurovascular networks that support the development and maintenance of thermogenic adipose tissue.

## RESULTS

### Formation of functional adipose depots from thermogenic versus non-thermogenic adipocyte

To generate cells for implantation we derived mesenchymal progenitors from human subcutaneous adipose tissue [21] as detailed in Materials and Methods, and cell suspensions were injected subcutaneously into each flank of nude mice (Extended Figure 1 A, B). Discernible adipose tissue pads, and circulating human adiponectin, were detected in all mice (Extended Figure 1 C, D), demonstrating that human adipocytes were able to secrete characteristic adipokines for an extended period after implantation. To monitor the development of adipocytes within the in vivo environment, we performed immunohistochemistry of the excised tissue with an antibody to perilipin 1 (PLIN1), which delineates lipid droplets. This analysis revealed a progressive increase in lipid droplet size over a period of 16 weeks (Fig. 1A). Notably, implantedadipocytes grew larger than those in endogenous mouse subcutaneous white adipose tissue as early as week4 after implantation and continued to increase to match the mean size of adipocytes in the human tissue from which progenitor cells were obtained (Fig. 1A, B). The change in mean adipocyte size was mostly attributable to a shift from multilocular to unilocular adipocytes (Fig. 1C). Expression of human *PLIN1* gene was relatively stable from weeks 4 to 16 (Fig. 1D, top panels), but *ADIPOQ* expression decreased as adipocytesize increased (Fig 1D, lower panels), mimicking the known inverse correlation seen in humans between ADIPOQ production and adipocyte hypertrophy [24]. Importantly, there was no detectableexpression of mouse *Plin1* or *Adipoq*, (Fig. 1D, top and lower panels), indicating that adipocytes in the developed depot are exclusively of human origin. These results indicate that as adipocytes continue to develop in the mouse they retain fundamental features of human adipose tissue, reflected by adipocyte sizeand regulated adiponectin expression.

**Figure 1.**
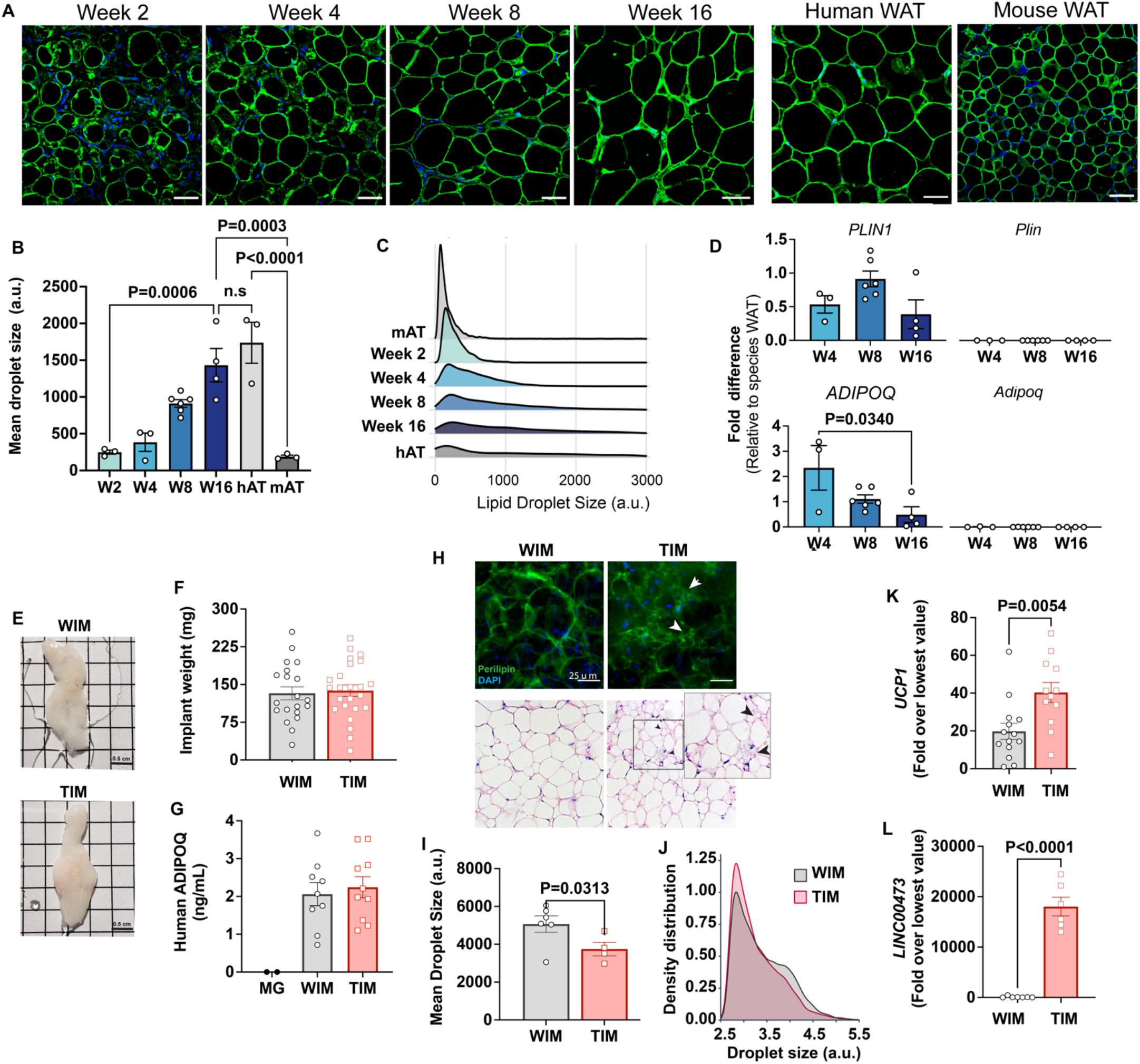
Characteristics of adipocytes during development of WIM and TIM. **A.** Representative images of PLIN1 (green) and DAPI (blue) staining of WIM at the indicated weeks after implantation, and of inguinal mouse and subcutaneous human adipose tissue, scale bar = 50 µm. **B.** Quantification of dropletsize of PLIN1-stained implants and tissues. Each symbol represents the mean area of all cells within three random fields of a single implant or tissue section. Bars represent the means and SEM of n=3 n=3, n=6, n=4, implants at weeks 2, 4, 8 and 16 respectively, and n=3 for independent human or mouse adipose tissue samples. Statistical significance of the differences between samples was calculated using one-way ANOVA with Tukey’s adjustment for multiple comparisons, and exact P-values are shown. **C.** Density distribution plot of droplet size of developing implants and control tissue. Each density distribution represents the combined density of all samples for each group (n = 3-6). **D.** RT-PCR of excised implants using species-specific probes for genes indicated. Values were normalized to those obtained from human or mouse adipose tissue samples exemplified in A. Symbols are the means of technical duplicates for each implant, and bars are the means and SEM of n=3, n=6 and n=4 implants at weeks 4, 8 and 16 respectively. Statistical significance of the differences between samples was calculated using one-way ANOVA with Tukey’s adjustment for multiple comparisons, and exact P-values are shown. **E.** Representative images of white implants (WIM) and thermogenic implants (TIM) at 8 weeks of development, grid size = 0.5 cm. **F.** Weightin mg of each excised implant from bi-laterally implanted mice. Bars are the mean and SEM of n=20 and n=24 implants for WIM and TIM, respectively. **G.** Circulating human-specific adiponectin (ng/ml) from bilaterally implanted mice. Symbols are the mean of technical duplicates for n=2, n=9 and n=10 mice implanted with Matrigel alone (MG), WIM or TIM respectively. **H.** Whole mounts of WIM and TIM at 8-weeks after implantation stained for PLIN1 (top panels), or with H&E. Arrowheads highlight clusters of dense, multilocular droplets in TIM. **I.** Mean droplet size (arbitrary units) in WIM and TIM. Symbol represent the mean areas of all cells from two independent sections from each implant. Bars represent the mean and SEM of n=6 and n=4 WIM and TIM implants, respectively. Statistical significance of the difference was calculated using unpaired, one-tailed Student t-test and resulting P-value is shown. **J.** Density distribution plot of droplet size of WIM and TIM. Each density distribution represents the averagedensity of all samples for each group. **K, L.** RT-PCR of excised implants using human-specific probes for the genes indicated. Symbols are the means of technical duplicates for each implant, and bars are the meansand SEM of n=14 and n=12 for *UCP1*, and n=7 and n=6 for *LINC00473,* in WIM and TIM, respectively.Values were normalized to the lowest expression level in the entire data set of n=26 implants. Statistical significance of the differences was calculated using unpaired, two-tailed Student t-tests, and exact P-valuesare shown.

We then asked whether depots formed from white (**W**hite **IM**planted=**WIM**) or thermogenically-induced adipocytes (**T**hermogenic **IM**planted=**TIM**) would be distinguishable from each other. The macroscopic features and weight of TIM and WIM were similar (Fig. 1E, F), as was the average concentration of circulating human adiponectin in mice harboring TIM compared to WIM (Fig. 1G). However, multilocularadipocytes could be detected in wholemount staining of TIM, by either PLIN1 immunostaining of whole mounts (Figure 1H, top panels, white arrows) or by H&E staining of tissue sections (Fig. 1H, lower panels, black arrows). Droplet quantification revealed a significant difference in mean droplet size (Fig. 1I), resulting from a higher density of small droplets (Fig. 1J). Expression of human *UCP1* was significantly higher in TIM (Fig. 1K), and expression of *LINC00473*, a marker for human thermogenic adipocytes [25],was exclusively detected in TIM (Fig. 1L). These results indicate that thermogenic features are maintainedfor a minimum of 8 weeks in adipose tissue developed from implanted cells.

### Beige adipocytes induce development of adipose tissue with a denser neurovascular architecture

To analyze the development of vasculature we stained whole-mounts and thin sections from the excised implants with endothelial stain isolectin (Fig. 2A). We detected dispersed endothelial cells as early as week2 after implantation in whole-mount specimens (Figure 2A, top panels). Between weeks 4 and 8, a well-developed vascular network, closely resembling that seen in human adipose tissue, was established. Vessels formed in apposition to enlarging adipocytes, as seen in sections co-stained with isolectin and antibodies to PLIN1 (Fig. 2A, lower panels). Quantification of the total intensity of isolectin staining in the thin sections, relative to PLIN1 staining to account for adipocyte mass, did not differ in a statistically significant way over time (Fig. 2B). However, quantification of the mean size of vascular structures stained by isolectin revealed a significant increase between weeks 4 and 8 (Fig. 2C), corresponding with the appearance of discernible vessels. Consistent with the intensity of isolectin staining, the expression of mouse vascular endothelial cadherin (*Cdh5*) did not differ in a statistically significant way over time and was exclusively of mouse origin as human *CDH5* was undetectable (Fig. 2D). These results suggest that endothelial cells invade the emerging tissue early in development, and subsequently mature into morphologically discernablevessels. Even though all vasculature was of mouse origin, the vascular network was morphologicallyindistinguishable from human WAT (Fig 2A, comparing Week 16 with human adipose tissue (Human AT)).

**Figure 2.**
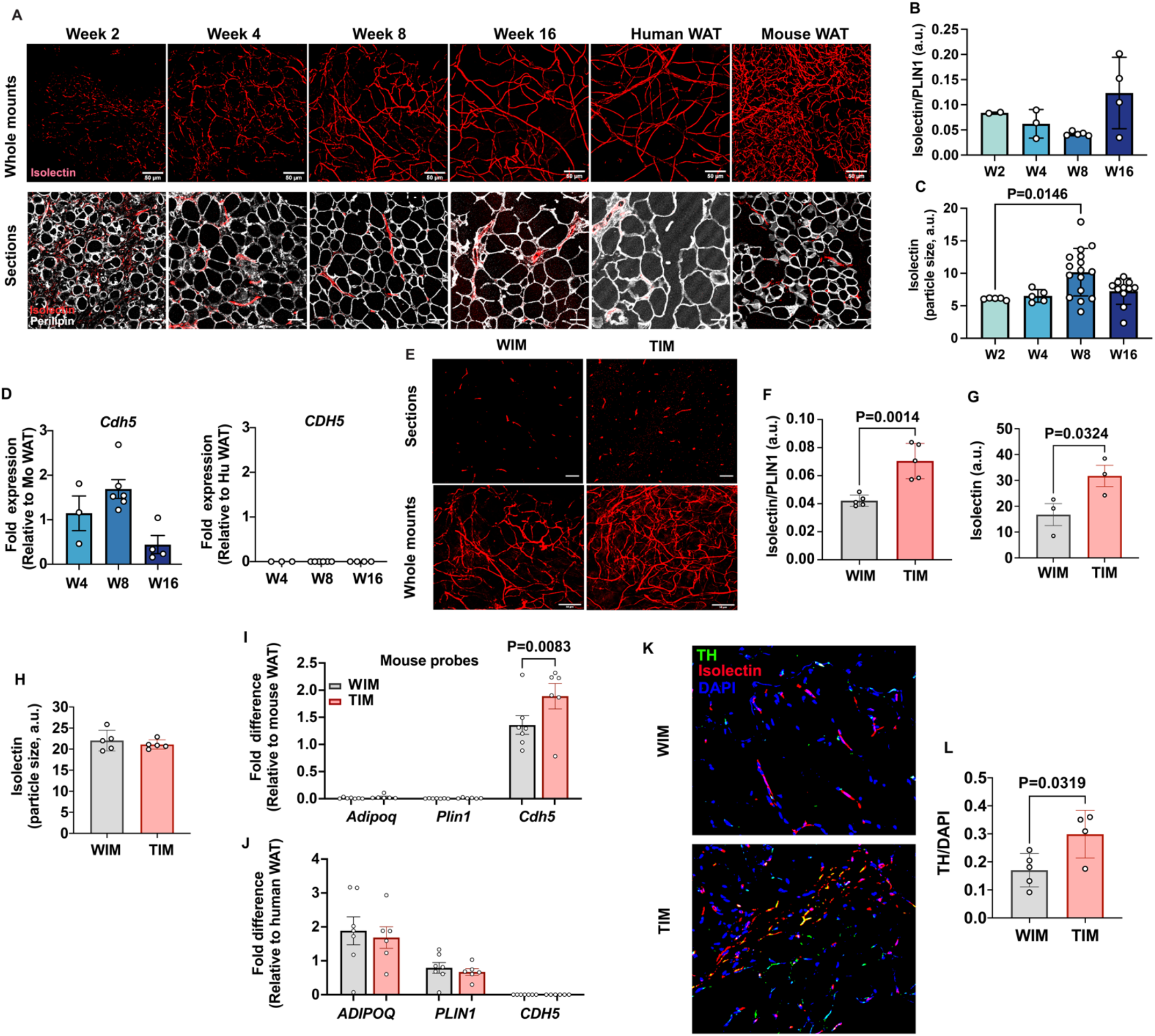
Vascular development of WIM and TIM. **A.** Representative images of isolectin staining of WIM at the indicated weeks after implantation, and of inguinal mouse and subcutaneous human adipose tissue, scale bar = 50 µm. **B.** Total isolectin staining in implants. Each symbol represents the sum of the areas of particles stained by isolectin, normalized to the total area stained by Plin1, of three random fields from a single implant. Bars represent the means and SEM of n=2, n=3, n=5, n=4, implants at weeks 2, 4, 8and 16 respectively. C. Mean size of isolectin-stained particles. Each symbol represents the mean area of particles stained by isolectin in three random fields per implant. Bars represent the means and SEM of n=5, n=5, n=16, n=11, implants at weeks 2, 4, 8 and 16 respectively. Significance of the differences between samples was calculated using one-way ANOVA with Tukey’s adjustment for multiple comparisons, and exact P-values are shown. **D.** RT-PCR of excised implants using species-specific probes for *CDH5*/*Cdh5*. Values were normalized to those obtained from mouse adipose tissue samples. Symbols are the means of technical duplicates for each implant, and bars are the means and SEM of n=3, n=6 and n=4 implants at weeks 4, 8 and 16 respectively. **E.** Representative images of thin sections (upper panels) and whole mounts(lower panels) of WIM and TIM after 8 weeks of implantation, stained with isolectin. **F, G.** Total isolectinstaining of thin sections and whole mounts, assessed as in (B). Bars represent the means and SEM of n=5, implants for thin sections, and n=3 for whole mounts of WIM and IM, respectively. Significance of the differences between samples was calculated using unpaired two-tailed Student t-tests. **H.** Mean size of isolectin-stained particles in WIM and TIM, analyzed as in (C). Bars represent the means and SEM of n=5,implants. **I, J.** RT-PCR of excised implants using mouse-specific (I) and human-specific (J) probes for the genes indicated. Values were normalized to those obtained from mouse and human adipose tissue samples.Symbols are the means of technical duplicates for each implant, and bars are the means and SEM of n=7, and n=6 WIN and TIM implants respectively. Significance of the differences between WIM and TIM for each gene was calculated using one-way ANOVA with the Holm-Sidak adjustment for multiple comparisons, and exact P-values are shown. **K.** Representative images of thin sections WIM and TIM after8 weeks of implantation, stained for isolectin (red), tyrosine hydroxylase (TH, green) and DAPI (blue). **L**.Total TH staining, where bars represent the means and SEM of n=5, and n=4 sections of WIM and TIM, respectively. Significance of the differences between samples was calculated using un-paired 2-tailed Student t-test and the exact P-value is shown.

We then compared the development of tissue vasculature between TIM and WIM after 8 weeks of implantation (Fig. 2E). Total intensity of isolectin staining in thin sections, relative to adipocyte mass as estimated by PLIN1 staining, was increased in TIM (Fig. 2F), as was the non-normalized intensity of isolectin in whole-mount specimens (Fig. 2G). The overall morphology of vascular structures was not different between WIM and TIM (Fig. 2H), suggesting that vessel maturation was unaffected. Gene expression levels of *Cdh5* were higher in TIM (Fig 2I), but those of human adipocyte genes *ADIPOQ* or *PLIN1*, were not different (Fig. 2J). These results indicate that thermogenic adipocytes induce the formation of a denser vascular network compared to non-thermogenic cells. We next asked whether TIM would havehigher content of blood cells, which we would expect if the newly formed vascular network was functional. For this we measured content of blood cells in the developing tissue, using the hematopoietic cell marker CD45 and the macrophage marker F4/80. We find that TIM had significantly higher number of macrophages compared to WIM (Extended Figure 2), consistent with a denser, functional vascular network.

Sympathetic innervation often accompanies vascularization [26–28]. To determine whether beige adipocytes also promote enhanced sympathetic nerve recruitment, we co-stained implants to simultaneously detect tyrosine hydroxylase (TH) and isolectin. Punctate staining consistent with nerve terminals was detected in all implants, with some overlapping with isolectin (Fig. 2K). Quantification revealed significantly higher TH staining in TIM compared to WIM (Fig. 2L), indicating enhanced sympathetic recruitment in conjunction with enhanced vascularization.

### The neurovascular network differentially responds to implanted adipocytes and supports TIM thermogenic phenotype

To determine the extent to which the neurovascular network elicited by thermogenic adipocytes contributes to the maintenance of their phenotype, we examined the autonomous capacity of adipocytes to retain thermogenic gene expression. We compared adipocytes that were never stimulated to adipocytes that were chronically stimulated with Forskolin (Fsk), and with adipocytes from which Fsk was withdrawn after a period of chronic stimulation (Fig. 3A). Removal of Fsk from chronically stimulated adipocytes resulted in a rapid increase in droplet size, which reached the size seen in cells that were never exposed to Fsk withindays of removal (Fig. 3B, C). Forskolin withdrawal also resulted in a sharp decrease in *UCP1*(Fig 3D) and *LINC00473* (Fig. 3E) expression, with values becoming statistically non-different from non-induced adipocytes within 5 days of withdrawal. Interestingly, the expression of *LINC00473* decreased even in the presence of chronic Fsk stimulation, while *UCP1* levels continued to increase over time, revealing different pathways of adaptation to chronic stimulation. Irrespective of these gene-specific differences, these results indicate that the thermogenic phenotype of adipocytes in-vitro is completely reversed within days of withdrawal of stimulation. We conclude that adipocytes lack the capacity to autonomously maintain a thermogenic phenotype, and that vascularization and innervation generated following implantation are likely to play a critical role in sustaining their thermogenic properties.

**Figure 3.**
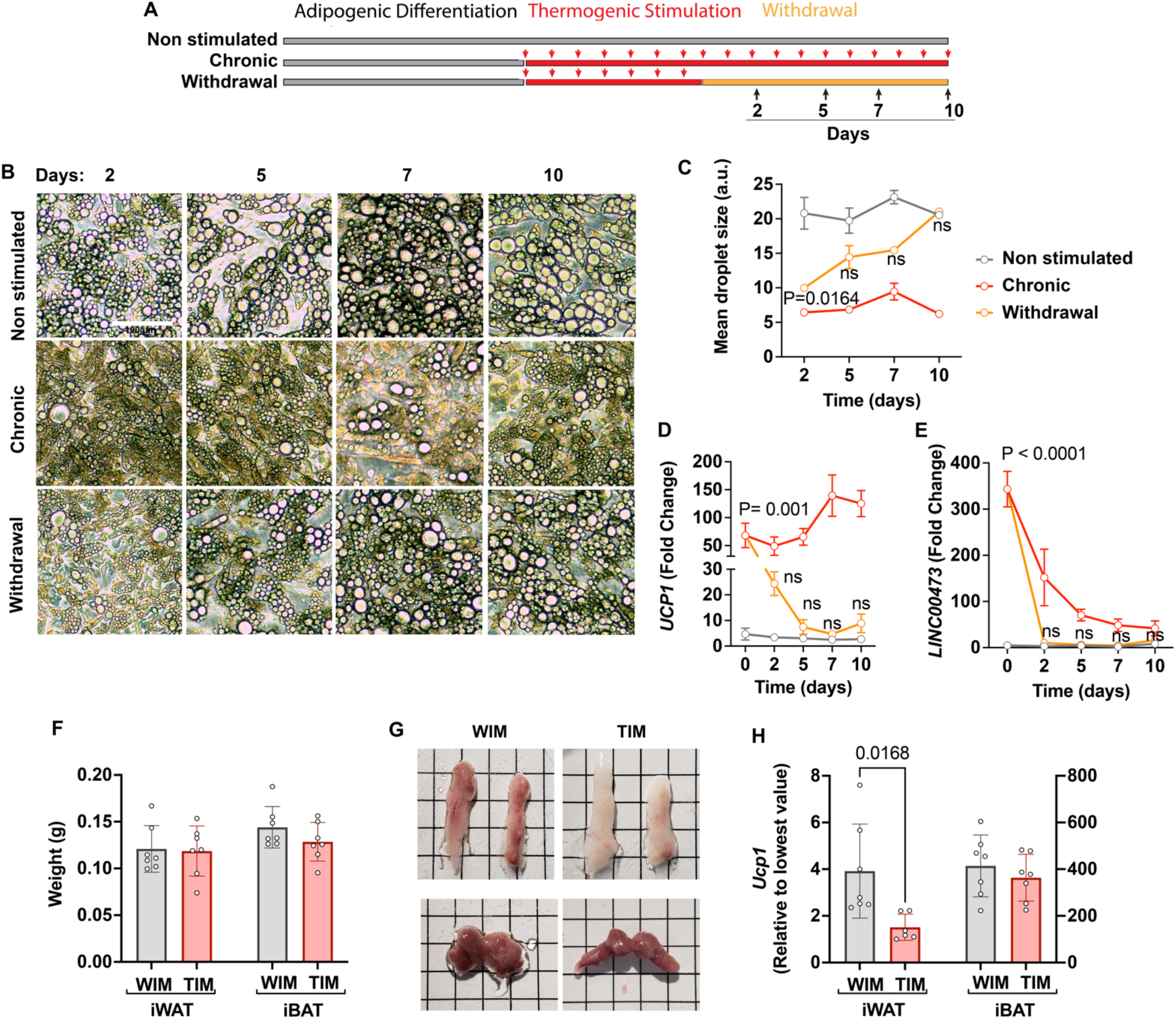
Capacity of thermogenic adipocytes to autonomously maintain their phenotype, and their effect on host adipose depots. **A.** Experimental paradigm for in vitro experiments. **B**. Phase images of cultured adipocytes during chronic stimulation and withdrawal. **C.** Quantification of droplet size, where symbols are the mean and SEM of all droplets in single images from each of n=3 independent culture wells.**D, E.** RT-PCR with human-specific probes for UCP1 (D) and LINC00473 (E), where symbols are the meanand error bars the SEM of n=3 independent culture wells assayed in technical duplicate. Values are expressed as the fold difference relative to the lowest value in the dataset. For (C), (D) and (E), statistical significance of differences at each time point between the withdrawal and the non-induced group, was measured using 2-way ANOVA with Dunnet’s multiple comparison test, and exact P-values are shown. **F.** Weight of excised inguinal (iWAT) and interscapular (iBAT) after 8 weeks of implantation with WIM or TIM. Symbols correspond to the summed weight of two inguinal or the single interscapular pad per mouse and bars the mean and SEM of n=7 mice. **G.** Representative images of excised fat pads. Top panels, iWAT;bottom panels iBAT. **H.** RT-PCR of *Ucp1*, where each symbol is the mean of technical duplicates, and bars represent the mean and SEM of n=6-7 pads. Data are expressed as the fold difference relative to the lowest value in the set, with scales for iWAT and iBAT on the left and right Y axes, respectively. Statisticalsignificance of differences was measured using non-paired, 2-tailed Student t-tests and exact P-values are shown.

Conversely, the interactions between implanted adipocytes and the neuro-vasculature might have systemic effects. To examine this possibility, we examined host adipose depots. The weights of mouse inguinal white and interscapular brown adipose tissue (iWAT and iBAT, respectively) were similar between mice implanted with either WIM or TIM (Fig. 3F), but the iWAT, which in athymic nude mice is noticeably beige, looked ‘whiter’ in mice harboring TIM compared to WIM (Fig. 3G). This morphological differencecorrelated with lower *Ucp1* expression in iWAT of TIM-implanted mice, while no differences were seen in iBAT (Fig. 3H). These results suggest that newly formed hybrid depots participate in feedback mechanisms that regulate total body content of thermogenic adipose tissue [29], potentially through their neurovascular architecture [30].

### Distinct gene expression features of adipocytes and host cells in WIM and TIM

To explore underlying mechanisms by which thermogenic adipocytes elicit a different vascularization and innervation and control systemic thermogenic adipose tissue, we performed RNA sequencing on WIM and TIM after 8 weeks of implantation. To distinguish transcriptomes corresponding to either mouse or human cells, fastq files were aligned to both genomes, and resulting reads were classified as either of human or mouse origin using the XenofilteR R-package. We then used mouse transcriptomic data to infer the mouse cell types that comprise WIM and TIM (Fig 4A). As a framework we used single-cell datasets from Burl etal. [31], who profiled stromovascular cells from both inguinal and gonadal mouse depots. Consistent with their findings, we distinguish 9 clusters, corresponding to adipose stem cells at distinct stages of differentiation, vascular endothelial cells, vascular smooth muscle cells, lymphocytes, dendritic cells, macrophages, neutrophils, and an unidentified cluster (Fig 4B). The most prominent host-derived cell populations in WIM and TIM, predicted by dampened weighted least squares deconvolution (DWLS, https://github.com/dtsoucas/DWLS), were adipose stem/progenitor cells, macrophages, and vascular endothelial cells, with vascular smooth muscle cells, dendritic cells, and other immune cells at very low numbers. No detectable differences in the relative proportion of cell types were observed between WIM and TIM (Fig 4C). It is interesting to note that progenitor cells accompanying vascularization of the developing tissue did not differentiate into adipocytes, as the expression of mouse *Plin1* and *Adipoq* in WIM and TIM is negligible (Fig. 2I).

**Figure 4.**
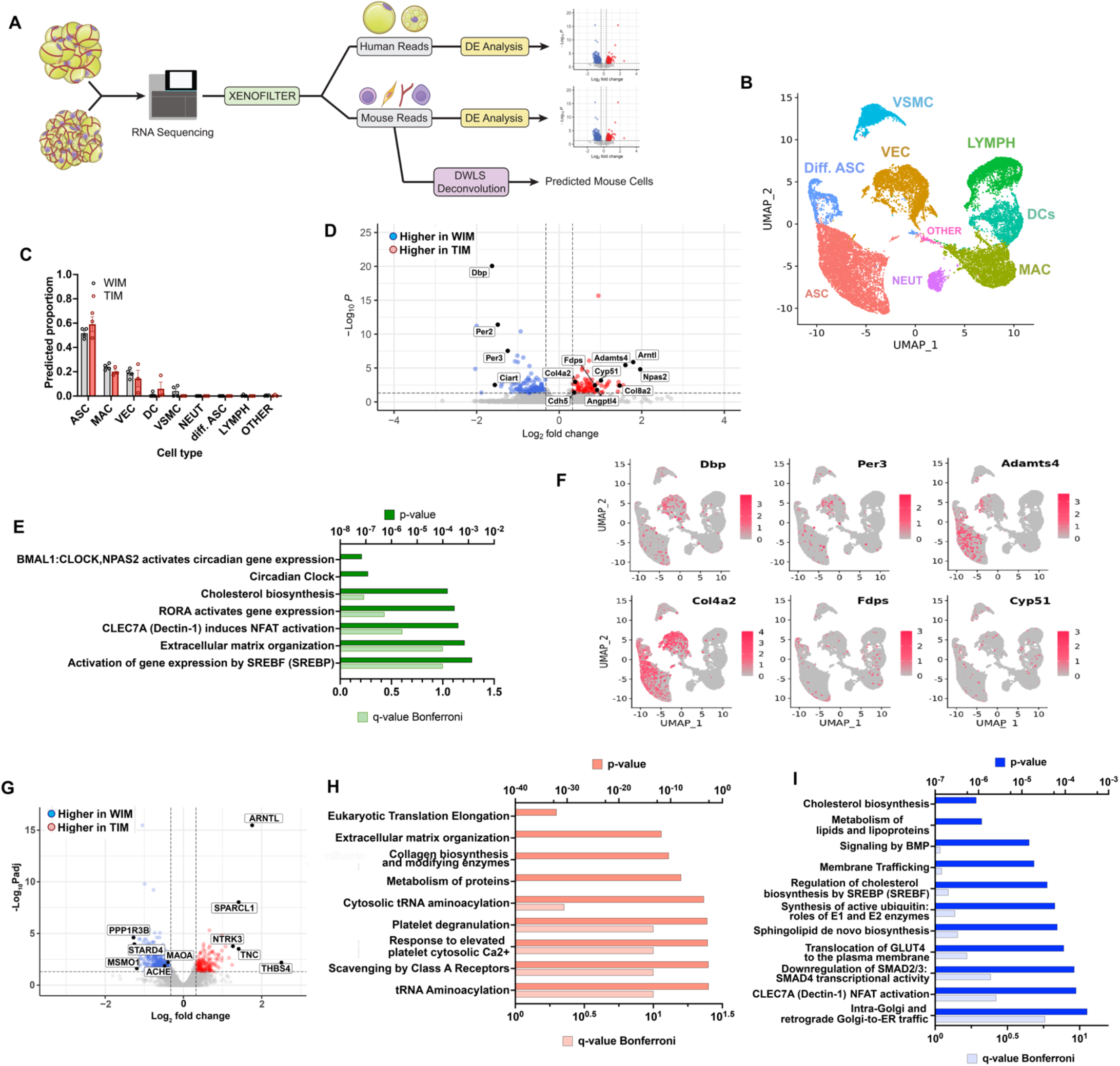
Distinct gene expression signatures of adipocytes and stromal cells in WIM and TIM. **A.** Overview of analysis paradigm. **B.** Dimensional reduction plot of data from Burl et al. [28], using UniformManifold Approximation and Projection (UMAP), where ASC = adipose stem cells, Diff. ASC = differentiating adipose stem cells, VEC = vascular endothelial cells, VSMC = Vascular smooth muscle cells, LYMPH = Lymphocytes, DCs = Dendritic cells, MAC = macrophages, NEUT = Neutrophils, OTHER = unidentified cluster. **C.** Predicted proportion of cells present in WIM and TIM. Symbols correspond to data from an individual implant and bars the mean and SEM of n=4 and n=3 WIM and TIM respectively. **D.** Volcano plot of genes mapping to the mouse genome which are differentially expressed between WIM and TIM. **E.** Pathway (REACTOME) enrichment analysis of mouse genes differentially expressed between WIM and TIM. **F.** Mouse genes differentially expressed between WIM and TIM mapped onto the UMAP representation shown in (A), with cells expressing the indicated genes highlighted in red. **G.** Volcano plot of genes mapping to the human genome differentially expressed between WIM and TIM. **H,I.** Pathway (REACTOME) enrichment analyses of human genes more highly expressed in TIM (H) or WIM (I).

We then probed for expression level differences in mouse genes between WIM and TIM. Of 13,036 detected genes, 86 were higher in TIM (range 1.25- to 414-fold), and 124 genes were higher in WIM (range 1.25- to8-fold), with a p-adjusted value lower than 0.05 (Fig 4D and Supplementary Table 1). Pathway analysis of differentially expressed genes revealed circadian clock genes as the most prominent pathway, with higher expression in TIM of *Npas2* and *Arntl* (3.9- and 3.5-fold higher in TIM, padj values 9.39E-05 and 7.36E-06 respectively) and higher expression in WIM of *Dbp*, *Ciart*, *Per2* and *Per3* (3.1-, 2.9-, 2.8- and 2.4-fold, padj values 1.28E-20, 0.008, 1.22E-11 and 4.99E-08 respectively). We also find enrichment in cholesterol biosynthetic and extracellular matrix remodeling pathways (Fig. 4E). We then asked if expressed genes correspond to a specific cell type, by searching for their expression in stromovascular cells from both inguinal and gonadal mouse depots shown in Figure 4B. Some of the differentially expressed genes associated with circadian rhythms (e.g., *Dbp*, *Per3*) or extracellular matrix remodeling (e.g., *Adamts4*, *Col4a2*) could be detected within clusters corresponding to progenitor cells andvascular endothelial cells (Fig. 4F). Genes in the cholesterol biosynthesis pathway (*Fdps* and *Cyp51*) are also seen in clusters corresponding to dendritic cells and macrophages. These results suggest that progenitorand endothelial cells respond to the different signals emanating from thermogenic versus non-thermogenicadipocytes and adopt a specific functional phenotype.

To elucidate mechanisms underlying differences in vascularization and innervation, we searched for genes differentially expressed between WIMand TIM that mapped to the human genome. Of 9655 detected genes, 515 were higher in WIM and 218 were higher in TIM, with a p-adjusted value lower than 0.05 (Fig. 4G, Supplementary Table 2). The most highly differentially expressed genes were associated with neurogenesis, including higher expression in TIM of *THBS4*, *TNC*, *SPARCL1* and *NTRK3* (5.6-, 2.7-, 2.7- and 2.4- fold, padj-values 0.006, 0.0003, 9.3E-09, and 0.0001, respectively) [29–34]. Pathway analysis of all genes with higher expression in TIM (Fig 4H) revealed enrichment in ribosomal protein-encoding genes, suggesting higher protein synthesis activity, followed by pathways affecting extracellular matrix composition, including numerous collagens. Many of these genes have been implicated in the control of blood vessel development, and could underlie the increased vascular density seen in TIM. As expected, *UCP1* mean values were higher in TIM (18.2 and 42.3 in WIM and TIM, respectively), consistent with the significantly elevated *UCP1* expression seen by RT-PCR (Fig 1K). Genes expressed at higher levels in WIM were associated with cholesterol and lipid metabolism as well as insulin action (Fig 4I), consistent with a more prominent role for lipid storage pathways in non-thermogenic adipocytes as they develop in vivo.

To identify genes that might be most relevant to the maintenance of the thermogenic phenotype, we searched for genes differentially expressed between non-thermogenic and thermogenic adipocytes prior to implantation that remain differentially expressed after 8 weeks. 988 genes were more highly expressed in thermogenic adipocytes, and 729 more highly expressed in non-thermogenic adipocytes (>2-fold, p-adjusted values < 0.05; Fig. 5A, Supplementary Table 3). Genes were assigned to Gene Ontology (GO) categories using https://go.princeton.edu/cgi-bin/GOTermMapper, a tool for mapping granular GO annotations to a set of broader, high-level GO parent terms [32]. Genes in thermogenic adipocytes disproportionally mapped to GO categories associated with secretion (Fig. 5B) and were enriched in pathways of extracellular matrix organization and signaling through GPCRs and IGFs (Fig 5C). Genes thatwere up regulated in non-thermogenic adipocytes were enriched in lipid metabolism pathways and transcriptional control of adipogenesis (Fig 5D). Of the 988 genes that were more highly expressed in thermogenic adipocytes prior to implantation, 21 were more highly expressed in TIM compared to WIM after 8 weeks (Supplementary Table 4), and 6 of these (*MMP14*, *ADAM19*, *TNC*, *ELN*, *COL12A1* and *TGFB3*) are associated with extracellular matrix organization, which was the only pathway enriched by these genes (Fig. 5E). Of the 729 genes that were expressed at higher levels in non-thermogenic adipocytesprior to implantation, 28 remained higher in WIM compared to TIM (Supplementary Table 4). Pathway analysis of these genes revealed neurotransmitter clearance as the top enriched pathway, with 2 genes, *ACHE* and *MAOA*, being higher in WIM.

**Figure 5.**
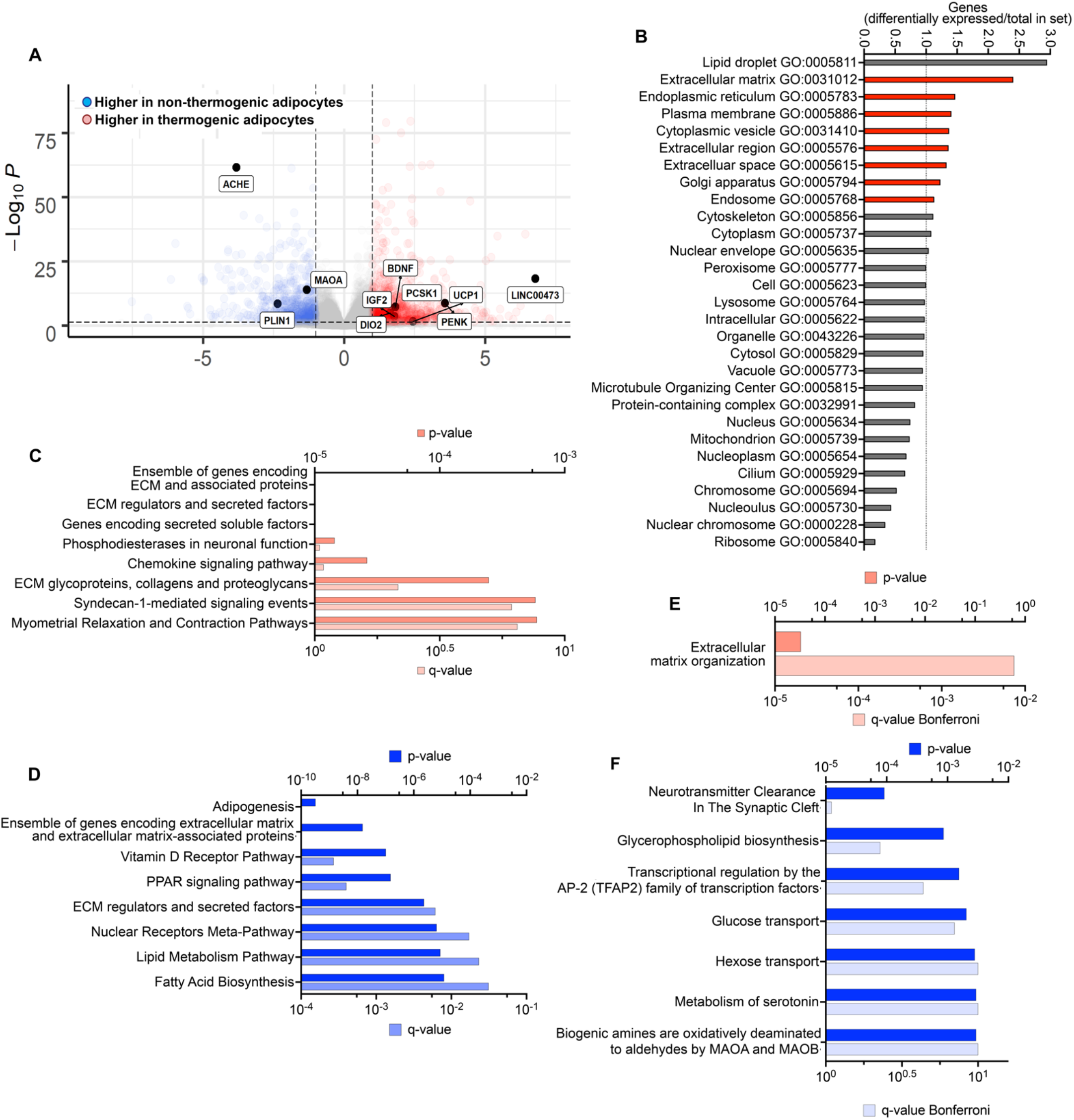
Differentially expressed genes persisting through tissue development. **A.** Volcano plot ofgenes differentially expressed between non-thermogenic and thermogenic adipocytes prior to implantation. Highlighted are selected genes characteristic of human thermogenic adipocytes and genes involved in the control of neurotransmission. **B.** Enrichment of GO categories by genes differentially expressed between non-thermogenic and thermogenic adipocytes, with categories for secretory proteins highlighted in red. **C, D.** Pathway (REACTOME) enrichment analyses of genes more highly expressed in thermogenic (C) or non-thermogenic (D) adipocytes. **E.** Pathway analysis of genes more highly expressed in thermogenic adipocytes that remained more highly expressed in TIM. **F.** Pathway analysis of genes more highly expressed in non-thermogenic adipocytes that remained more highly expressed in WIM.

A lower expression of *ACHE* and *MAOA* in adipocytes would be expected to enhance neurotransmitter tone and maintenance of the thermogenic phenotype. Indeed, in mouse models, Maoa activity has been implicated in the regulation of norepinephrine stimulated adipose tissue lipolysis and browning, but its expression has been localized to a specific macrophage population [33–35]. Our finding of *MAOA* transcripts in WIM and TIM suggested expression in human adipocytes. To confirm that *MAOA* transcriptsare indeed present in adipocytes in endogenous human adipose tissue, we re-analyzed single-nuclei RNA Seq data generated by Sun et. al. [36]. We reproduced their main findings, including the presence of a largeproportion of immune cells, and a clearly defined adipocyte population (Fig. 6A). *MAOA* transcripts were detected exclusively in the population of cells also expressing *ADIPOQ*, which correspond to adipocytes (Fig. 6B). Adipocyte, smooth muscle cells and fibroblast nuclei also contained transcripts for *SLC22A3*, themajor extra-neuronal catecholamine transporter, indicating that human adipocytes express both genes required for norepinephrine clearance (Fig 6B). In contrast, neither *Maoa* nor *Slc22a3* transcripts were detectable in mouse cells expressing *Adipoq* [31] (Figure 6C). Parallel RT-PCR and western blotting analysis of human adipocytes at different stages of differentiation in vitro indicate that transcriptional levels of *MAOA* reflect protein expression (Fig 6D). Moreover, MAOA protein is exclusively detected in adipocytes in human but not mouse adipose tissue (Fig 6E), and in adipocytes in WIM, but not in vicinal endogenous mouse adipocytes (Fig 6F), despite being detectable in mouse immune cells (Figure 6F).

**Figure 6.**
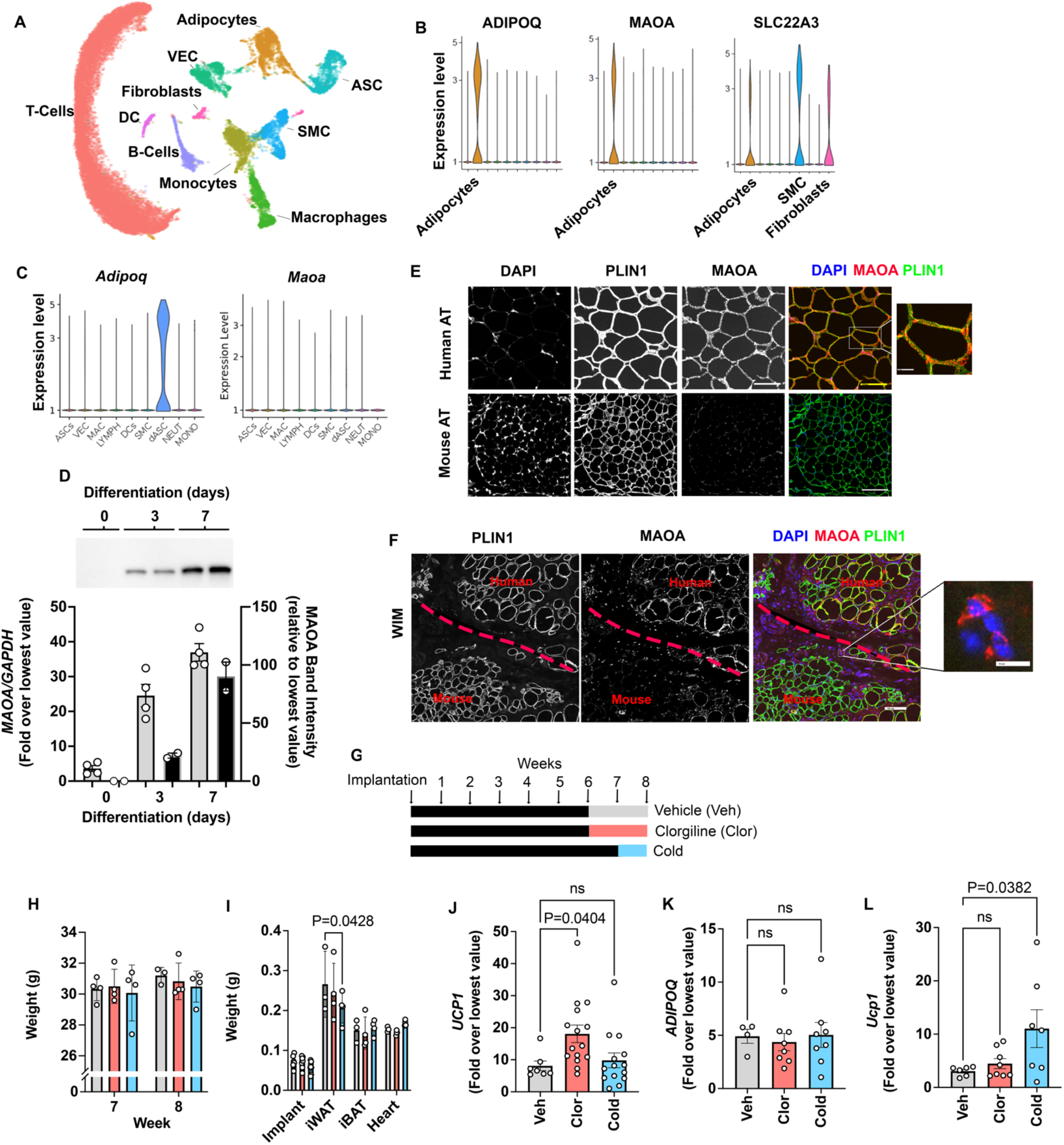
Species specific expression of MAOA in adipocytes. **A.** Dimensional reduction plot of datafrom Sun et. al. [39], using Uniform Manifold Approximation and Projection (UMAP). **B, C.** Violin plots of indicated human and mouse genes from single cell data from Sun et. al. [39] (B) and Burl et al. [28] (C). **D.** RT-PCR and western blotting at indicated days following induction of differentiation in human adipocytes. Samples were prepared from 2 independent wells and subjected to western blotting with an antibody to MAOA. The intensity of the single detected band was quantified, and values plotted in the right Y axis. RT-PCR results are plotted in the left Y axis. Symbols represent the fold difference over the lowest value in the dataset and bars the mean and SEM of n=4 independent cultures. **E, F.** Representative images of human and mouse subcutaneous adipose tissue (E) and of WIM and vicinal mouse adipose tissue (F), stained with antibodies to MAOA and PLIN1, and counterstained with DAPI. Bars = 100 um. Expanded section in (F) shows staining for MAOA in immune cells within mouse tissue. Bar= 10 um. G. Experimental time line for exposure on mice harboring WIM to clorgyline or cold. H. Weight of mice at 7 and 8 weeks I. Weights of dissected tissues at 8 weeks. J,K. RT-PCR for human *UCP1* (J) and *ADIPOQ* (K) in excised WIM. Bars represent the mean and S.E.M and symbols correspond to each independent implant (n=7-8 implants). L. RT-PCR for mouse *Ucp1* in excised inguinal adipose tissue (n=7-8 fat pads). Bars represent the mean and S.E.M and symbols correspond to each independent fat pad. H-L, Statistical significance of the differences was assessed using one way ANOVA with Dunett’s multiple comparison correction, and exact values are shown.

To determine whether expression of MAOA in human adipocytes is functionally relevant, mice harboring WIM were treated with a low dose of the specific MAOA inhibitor clorgyline for 2 weeks. A separate cohort of mice was gradually acclimated to low ambient temperature as an alternative method to enhance adrenergic tone (Fig 6G). Neither treatment resulted in significant changes in body weights (Fig 6H), nor in weights of most tissues with the exception of a decrease in inguinal subcutaneous adipose tissue weight in cold-acclimated mice (Fig 6I). However, implants from clorgyline-treated mice expressed significantly elevated *UCP1* (Figure 6J), with no change in *ADIPOQ* expression (Fig. 6K). Importantly, *Ucp1* in mouse subcutaneous adipose tissue was not affected by clorgyline, despite it being responsive to cold stimulation (Fig 6L), demonstrating that the effect of clorgyline was specific for tissue containing human adipocytes.

## DISCUSSION

Efforts to understand mechanisms of human thermogenic adipose tissue development are limited by the difficulty of accessing thermogenic adipose tissue from human subjects, and from paucity of models to study human tissue development. Here we leverage a species-hybrid model in which adipose tissue is formed after implantation of developing human adipocytes into mice. This model is enabled by methods to generate multipotent human progenitor cells that can be driven towards differentiation into multiple adipocyte subtypes [19]. Adipocytes complete maturation *in vivo*, developing key species-specific features, such as larger cell size compared to mouse adipocytes, and maintaining the thermogenic phenotype induced in-vitro. Developing adipocytes recruit additional host cellular components, including vascular endothelial cells, adipose stem/progenitor cells, immune cells and neuronal components to make functional adipose tissue.

An important question resolved by this work is whether the vasculature of thermogenic adipose tissue differs qualitatively from that of non-thermogenic tissue. By distinguishing adipocytes from other cellular components at the transcriptional level and complementing these data with single cell atlases [37–39], we find differences in vascular endothelial cells of thermogenic adipose tissue, including significant increase in *Col4A2*,which is associated with small vessel stability [38], and lower levels of circadian clock genes *Dbp*, *Per2* and *Per3*, which oscillate in adipose tissues of mice and humans [40–43], and have been associated with the activity of thermogenic adipocytes [44, 45]. Our finding that these genes are expressed in the vasculature of adipose tissue, rather than in the adipocytes, a finding that may explain the parallel oscillations of these genes in other peripheral tissues [46, 47].

The vasculature of the hybrid tissue may also be critical in defining its immune composition. We find that the relative proportion of immune cells in the hybrid tissue resembles that of mouse adipose tissue, which differs from human by having a relatively high macrophage content [48]. Cells within the vasculature, such as endothelial or progenitor cells, might be more important in defining the immune composition of adipose tissue than adipocytes [49, 50]. Nevertheless, the formation and features of the mature vascular network are initiated by developing adipocytes, as implanted cells lack vascular progenitors, and mouse endothelial cells infiltrate the tissue prior to the formation of blood vessels as seen in other systems [39]. Implanted adipocytes express high levels of secreted and extracellular matrix proteins, including collagens associated with angiogenesis. Further experiments will be required to delineate which of these genes are most relevant in establishing the adipose tissue vascular architecture and the distinct features associated with thermogenic adipose tissue.

In addition to a qualitative and quantitative different vasculature, thermogenic adipocytes also recruit a denser neuronal sympathetic network. The ability of implanted thermogenic adipocytes to recruit this neurovascular network is critical, as in the absence of external stimuli adipocytes lose their thermogenic phenotype within days. Indeed, at 8 weeks of development the most highly differentially expressed thermogenic adipocyte genes are associated with neurogenesis (*THBS4*, *TNC*, *NTRK3* and *SPARCL1*) [51–56]. Interestingly, two of these factors, Thbs4 and Sparcl1, have been identified as factors in blood from young mice that can directly stimulate synaptogenesis [51]. Beige adipocytes have been found to be neuroprotective [57], raising the possibility that these neurogenic factors produced by thermogenic adipocytes might not only enhance local innervation but might contribute to neuroprotection.

In addition to enhanced levels of neurogenic factors, we found that thermogenic adipocytes display decreased expression of neurotransmitter clearance enzymes MAOA and ACHE. MAOA catalyzes the oxidative deamination of norepinephrine and may affect thermogenesis by dampening adrenergic tone. ACHE is responsible for the catabolism of acetylcholine, which has been found to activate thermogenic adipocytes in the subcutaneous adipose tissueof mice [58]. Importantly Maoa has been implicated in the control of adipose tissue thermogenesis in mice [33, 34], through its expression in a specific subset of macrophages. Our studies indicate that in contrast to mice, human *MAOA* is strongly expressed in adipocytes, which also express the non-neuronal transporter *SLC22A3*, allowing for transport and degradation of catecholamines.

Our findings and those of others [59] suggest that adipocytes themselves play a major role in establishing adrenergic tone and controlling thermogenic activity in humans, with potential metabolic effects, as MAOA is increased in visceral adipose tissue of obese compared to non-obese patients [60], concomitantly with decreased expressionof UCP1 [61–63]. Our studies directly demonstrate that inhibition of MAOA with low doses of clorgyline induces expression of UCP1 in hybrid tissue, but not in mouse tissue, highlighting the functional relevance of the human adipocyte-expressed enzyme. The finding that inhibition of MAOA induces human adipocyte beiging in vivo suggests brain impenetrable, adipocyte favoring formulations of MAOA inhibitors as potential therapeutic agents in metabolic disease.

## MATERIALS AND METHODS

### Human Subjects

Abdominal adipose tissue was collected from discarded tissue of patients undergoing panniculectomy, with approval from the University of Massachusetts Institutional Review Board (IRB ID 14734_13).

### Generation of cells and implants

Small pieces of fat (∼1mm^3^) from the excised human adipose tissue were embedded in Matrigel (200 explants/10 cm dish) and cultured for 14 days as described before [21]. Single-cell suspensions were obtained using dispase and plated into 150 mm tissue culture dishes. After 72 h, cells were split 1:2, grown for an additional 72 h, recovered by trypsinization and frozen. To generate the implants, ∼10E7 cells were thawed into each 150 mm plate, and after 72 hours split 1:2 at a dense seeding density of ∼8E6 cells per 150 mm plate. Upon confluence (approximately 72 hours after plating), differentiation was induced by replacing the growth media with DMEM +10% FBS, 0.5 mM 3-isobutyl-1-methylxanthine, 1 µM dexamethasone, and 1 μg/mL insulin (MDI). MDI media was changed daily for 72 hours at which time the differentiation medium was replaced by DMEM + 10%FBS, 50% of which was replaced every 48 hours for 10 days. After 10 days of differentiation, a subset of cells was stimulated daily for 7 days with forskolin (1 µM final concentration) to induce the thermogenic phenotype. Single cell suspensions of differentiated and thermogenic-induced adipocytes were obtained by incubation for 7-10 min in Trypsin (1X)/collagenase (0.5 mg/mL). Proteases were quenched by dilution into culture media, cells pelleted by centrifugation at 500 rpm for 10 minutes. The media layer between floating and pelleted cells was removed, and remaining cells brought to 1 mL total volume with ice-cold PBS, placed on ice andmixed with an equal volume of ice-cold Matrigel. 0.5 mL aliquots of the cell suspension were placed in 1 ml syringes on ice and injected subcutaneously into each flank of immunodeficiency male nude mice (Nu/JJackson labs stock no: 002019) using a 20G needle. At each experimental time point, developed implants were dissected and snap frozen until further analysis.

### Human-specific adiponectin ELISA

Blood was obtained through cardiac puncture, centrifuged, and the plasma was collected and stored at -80°C. A human-specific adiponectin ELISA (Invitrogen KHP0041) wasused to measure the concentration of human adiponectin in mouse blood.

### RNA extraction and quantitative PCR

Implants were placed on TRIzol (Invitrogen) and homogenized with Tissuelyser (Qiagen). Total RNA was reverse transcribed using the iScript™ cDNA Synthesis Kit (BioRad) per manufacturer’s protocol. Either SYBR Green or PrimePCR™ Probes (BioRad) were used todetect amplification of human- and mouse-specific genes using the following primers:

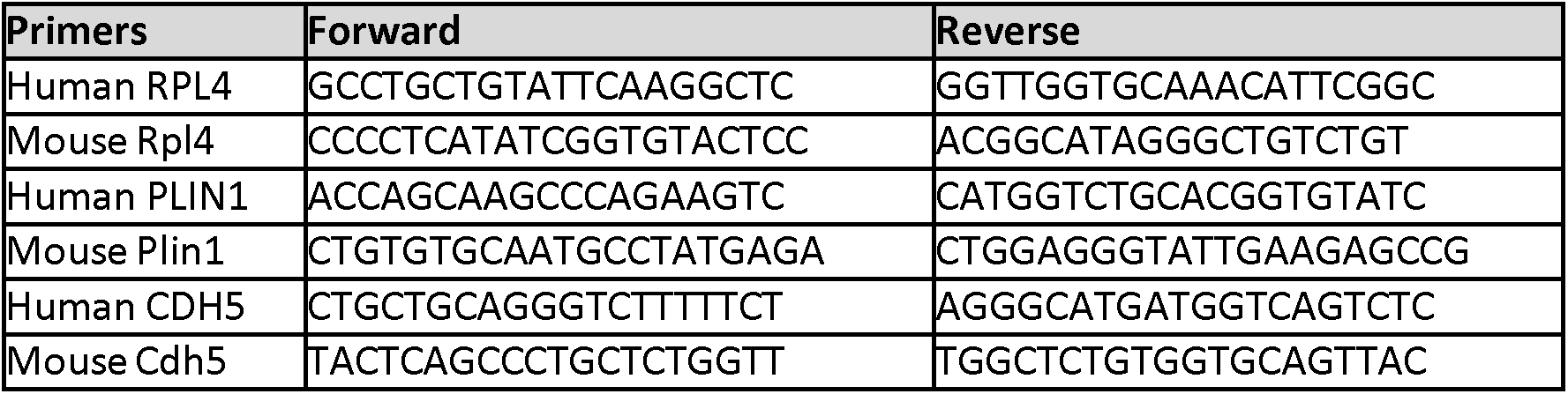

### RNA-Seq

RNA was extracted from white and thermogenic implants using the TRIzol method. Library preparation was performed using the TruSeq cDNA library construction (Illumina). Samples were processed on the Illumina HiSeq 550 sequencing system with the NextSeq 500/550 High Output kit v2.5 (Illumina, Cat. No. 20024906). The generated fastq files were loaded into the DolphinNext platform (https://dolphinnext.umassmed.edu/) and the Bulk RNA sequencing pipeline was used. The .fastq files were aligned to both the human (hg38) and the mouse (mm10) genome. The resulting alignments were processed using the R-package XenofilteR to classify reads as either of human or mouse origin. Reference (https://github.com/PeeperLab/XenofilteR) for more details on XenofilteR source code. Once aligned, the files were run through RSEM for normalization. Differential expression analysis was performed using the DEBrowser platform (https://debrowser.umassmed.edu/). Gene ontology analysis was performed by combining results from TopFunn (https://toppgene.cchmc.org/).

### Single Cell RNA-Seq Processing

Single-cell RNA sequencing data was obtained from Burl, Rayanne B.,et al. 2018. The provided .fastq files were loaded into DolphinNext and processed using the Cell Ranger pipeline. The output files from Cell Ranger were then loaded into the Seurat(https://satijalab.org/seurat/index.html). We removed unwanted cells from the data by sub-setting features as follow: nFeature_RNA > 100 & nFeature_RNA < 5000. We then proceeded to normalize the data usinga logarithmic normalization method with a scale factor of 10000. We then applied a linear transformation to scale the data followed by dimension reduction analysis (PCA). Following PCA analysis we determined that the first 8 principal component analysis were optimal for further analysis. To cluster the cells, we performed non-linear dimension reduction analysis using the UMAP algorithm. Analysis was done using the first 8 principal components and a resolution of 0.3. Finally, differentially expressed genes for each cluster were identified to determine biomarkers for cell cluster identification.

### Cell Predictor Using DWLS Deconvolution

Source code for DWLS deconvolution can be found at https://github.com/dtsoucas/DWLS. Briefly, metadata from the Seurat object was used to generate a matrix containing the reads from the scRNA and one containing the cluster labels for each gene. A signature matrix was generated using the MAST function provided with the DWLS deconvolution code. Only genes in common between the scRNA and the bulk RNA dataset were used for prediction. DWLS deconvolution was performed using the solveDampenedWLS function and proportions of predicted cells were plotted.

### Single Nuclei RNA-Seq Processing

Seurat objects for single nuclei RNA sequencing was kindly providedby the Wolfrum lab, who published their analysis [36]. Briefly, the Seurat objects containing the raw data were loaded the Seurat package, and two different samples were processed: single nuclei from human brown adipose tissue, single nuclei from mouse brown adipocytes. Briefly, for the human brown adipose tissue dataset, we first removed unwanted cells from the data by sub-setting features as follow: nCount_RNA > 1000 & nCount_RNA < 8000 & prop. mito < 0.3. We then normalized the data using a logarithmic normalization with a scale factor of 10000. We followed this by applying a linear transformation to scale the data followed by dimension reduction analysis (PCA). We determined that the first 10 principal component analysis were optimal for further analysis. To cluster the cells, we performed non-linear dimension reduction analysis using the UMAP algorithm. Analysis was done using the first 8 principal components and a resolution of 0.3. For the mouse brown adipocytes, we first removed unwanted cells fromthe data by sub-setting features as follow: nFeature_RNA > 1000 & nFeature_RNA < 5000. We then normalized the data using a logarithmic normalization with a scale factor of 10000. We followed this by applying a linear transformation to scale the data followed by dimension reduction analysis (PCA). We determined that the first 8 principal component analysis were optimal for further analysis. To cluster the cells, we performed non-linear dimension reduction analysis using the UMAP algorithm. Analysis was doneusing the first 8 principal components and a resolution of 0.2. Finally, clusters names were assigned using SingleR (https://github.com/dviraran/SingleR). The human primary cell atlas [64] was used as a reference to identify clusters based on differentially expressed genes of human brown adipose tissue.

### Histochemistry and quantification

All samples were fixed in 4% paraformaldehyde overnight at 4°C and thoroughly washed with PBS. Tissue sections (8 µm) were mounted on Superfrost Plus microscope slides (Fisher Scientific) and stained with hematoxylin and eosin. For whole-mount staining, tissue fragments (∼1mm^3^) were stained and mounted between 1.5 mm coverslips sealed with ProLong Gold Antifade Reagent (Life Technologies). Mouse vasculature was detected using isolectin GS-IB4 AlexaFluor-647 conjugate (Invitrogen I32450) and human vasculature using Ulex Europaeus Agglutinin I (UEA I),DyLight® 594 (Vector Laboratories DL-1067-1). Nuclei were stained using DAPI (Life Sciences 62249). Perilipin-1 was detected with an antibody that recognizes both human and mouse protein (CellSignaling #9349). MAOA was detected using CellSignaling #73030 (human) or CellSignaling #75330 (mouse). To identify sympathetic nerves an antibody for Tyrosine Hydroxylase was used (Millipore Sigma - AB152). Macrophages were identified by co-staining using antibodies for CD45 (Abcam – ab8216) and F4/80 (Abcam – ab6640). Tissue sections were imaged using ZEISS Axio Scan Z1. Whole-mount images were acquired using an Olympus IX81 microscope (Center Valley, PA) with dual Andor Zyla sCMOS 4.2 cameras (Belfast, UK) mounted on an Andor TuCam two camera adapter (Belfast, UK).

Image analysis was performed using the open software platform FIJI. For all images to be compared, a single stack was generated and subjected to the same filtering and quantification processes. Adipocyte areawas determined from PLIN1 stained images imported as a single stack, converted to 8-bit grayscale, binarized, eroded twice, and object number and size measured through particle analysis (size = 350-2900 um, circularity = 0.25-1). Density distributions (scaled) were generated using R-studio through the density distribution function in ggplot2. Vascular density and size were measured using isolectin stained images, imported as a single stack into FIJI, background subtracted (rolling radius = 1) to enhance contrast, binarized using the Threshold function and subjected to particle analysis. The total vascular area of each section was the sum of the individual particles. The average vascular size was the mean of the individual particles. For macrophage quantification, CD45, F4/80 and DAPI images were background subtracted (rolling radius = 1) and binarized. To isolate macrophages that were both CD45 and F4/80 positive we usedthe image calculator function (image calculator > operation: AND). Particle analysis was performed, and the number of particles normalized to the number of nuclei present in the whole section to account for differences in section sizes. Sympathetic density was calculated using TH and isolectin stained images after background subtraction (rolling radius = 1) and binarization. Total sympathetic area in the section was thesum of all particles normalized to the number of nuclei present in the whole section to account for differences in section sizes.

### Thermogenic withdrawal of thermogenic adipocytes

Cells (∼6.0×10^6^) were thawed and seeded on 6-well plates, differentiated into white adipocytes, and stimulated with forskolin as described above. After chronic forskolin stimulation, a subset of cells was removed from daily stimulation for 10 days. Images were taken and samples were placed on TRIzol (Invitrogen) at days 0, 2, 5, 7, and 10 of withdrawal. Samples were homogenized using the Tissuelyser (QIAGEN), and total RNA was then isolated for analysis as described above.

## ACKNOWLEDGEMENTS

This study was supported by NIH grants DK089101 and DK123028 to SC and GM135751 to JSR. We acknowledge the use of services from the UMASS SCOPE core for high resolution confocal imaging, the biomedical imaging (BIG) facility for epifluorescence imaging and image analysis consultation. Finally, we acknowledge the use of services from the UMASS Morphology Core facility for sample preparation and sectioning for histological analysis.

## AUTHOR CONTRIBUTIONS

JSR: hypothesis generation, conceptual design, experiment design and performance, data analysis, manuscript preparation. ZYL: conceptual design, experiment design and performance, data analysis, manuscript preparation. TD: experiment design and performance, data analysis, manuscript preparation. AD: experiment design and performance, data analysis, manuscript preparation. QY: conceptual design, manuscript preparation. RRR: conceptual design and performance. PS: conceptual design and performance. SJ: conceptual design, manuscript preparation. DZ: conceptual design, manuscript preparation. TN: conceptual design, manuscript preparation. SC: supervision of work, hypothesis generation, conceptual design, manuscript preparation.

---

**Extended Figure 1.**
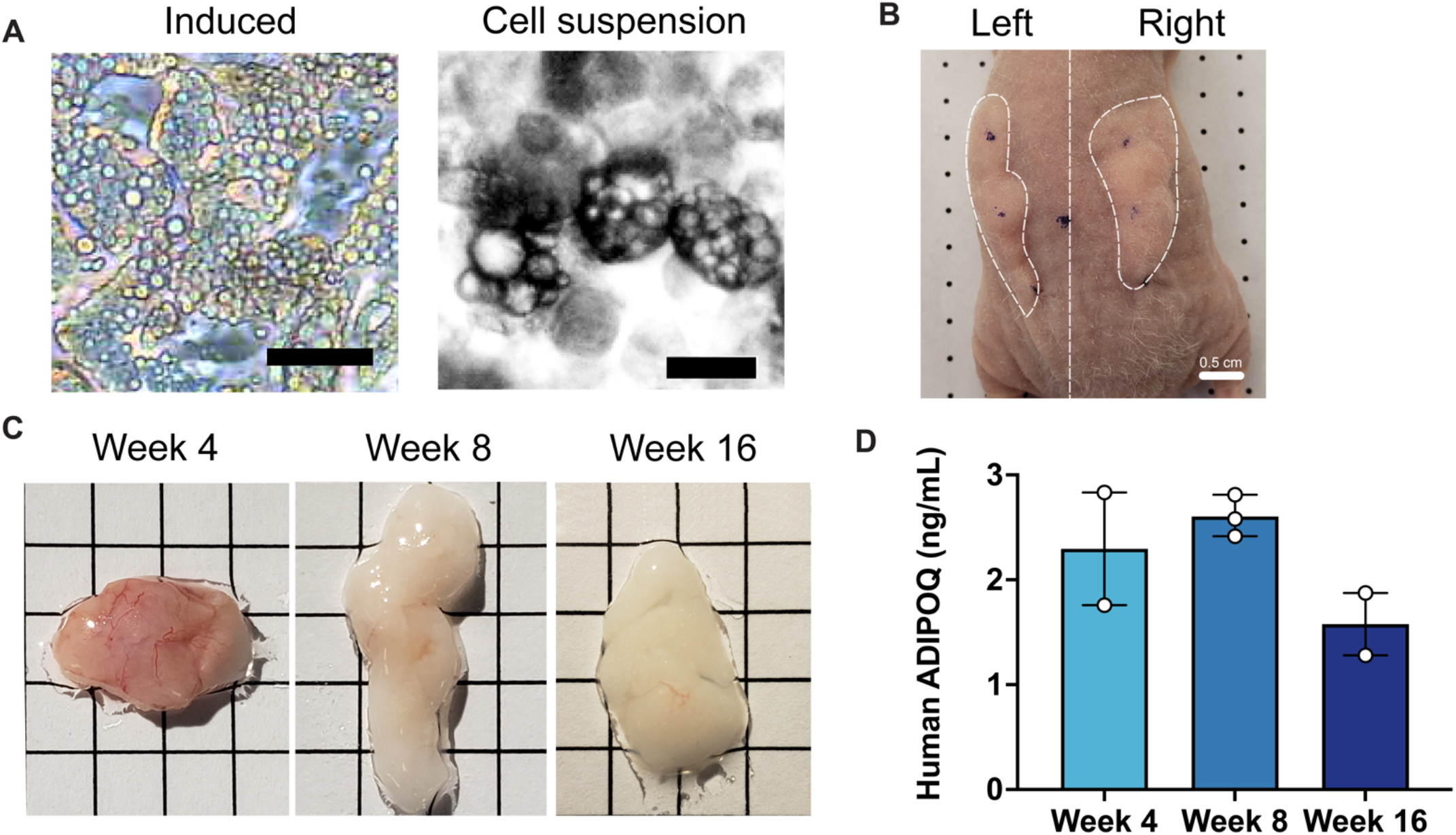
Development of implants from cultured adipocytes in mice. **A.** Representative images of differentiated adipocytes and single cell suspension used for implantation, scale bar = 25 µm. **B.** Representative image of bilateral flank implantation, scalebar = 0.5 cm. **C.** Excised implants at 4, 8, and 16 weeks of development, grid size = 0.5 cm. **D.** Circulating human adiponectin at indicated weeks after implantation. Symbols represent the mean of technical duplicates for each individual mouse, bars represent the mean and range of n=2, n=3 and n=2 mice for weeks 4, 8 and 16 respectively.

**Extended Figure 2.**
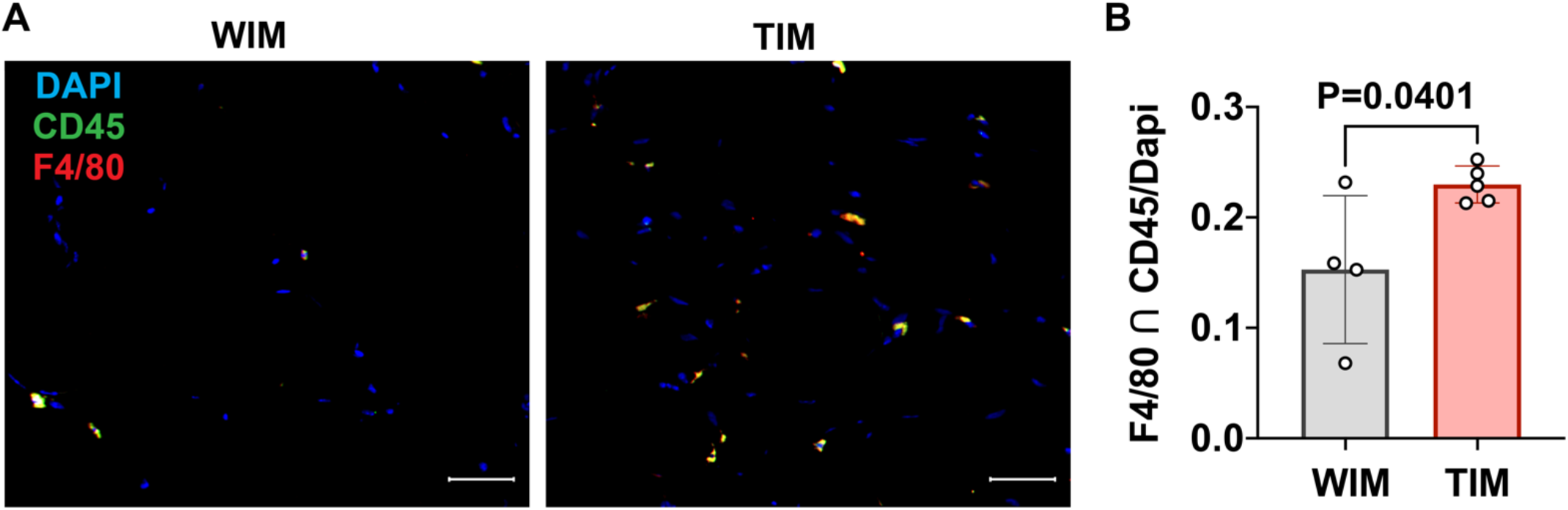
Density of macrophages in WIM and TIM. **A.** Representative images of thin sectionsWIM and TIM after 8 weeks of implantation, stained for F4/80 (red), CD45 (green) and DAPI (blue). **B**. Macrophage staining, where symbols are the mean of three independent sections per implant, and bars represent the means and SEM of n=4 and n=5, WIM and TIM implants, respectively. Data represents the cells stained with both CD45 and F4/80, normalized to DAPI to account for section size. Significance of the differences was calculated using un-paired 2-tailed Student t-test and the exact P-value is shown.

**TABLE 1:**
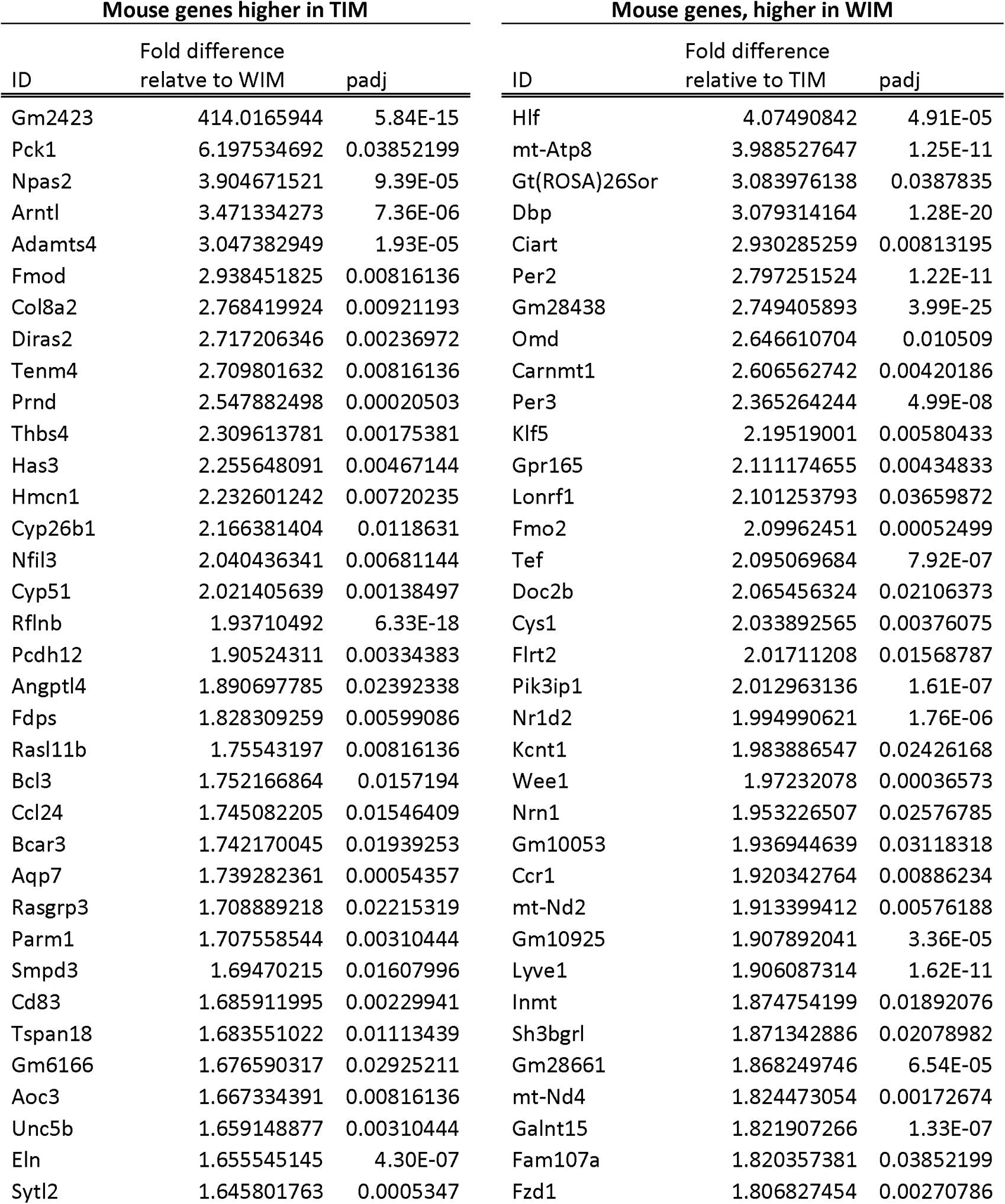
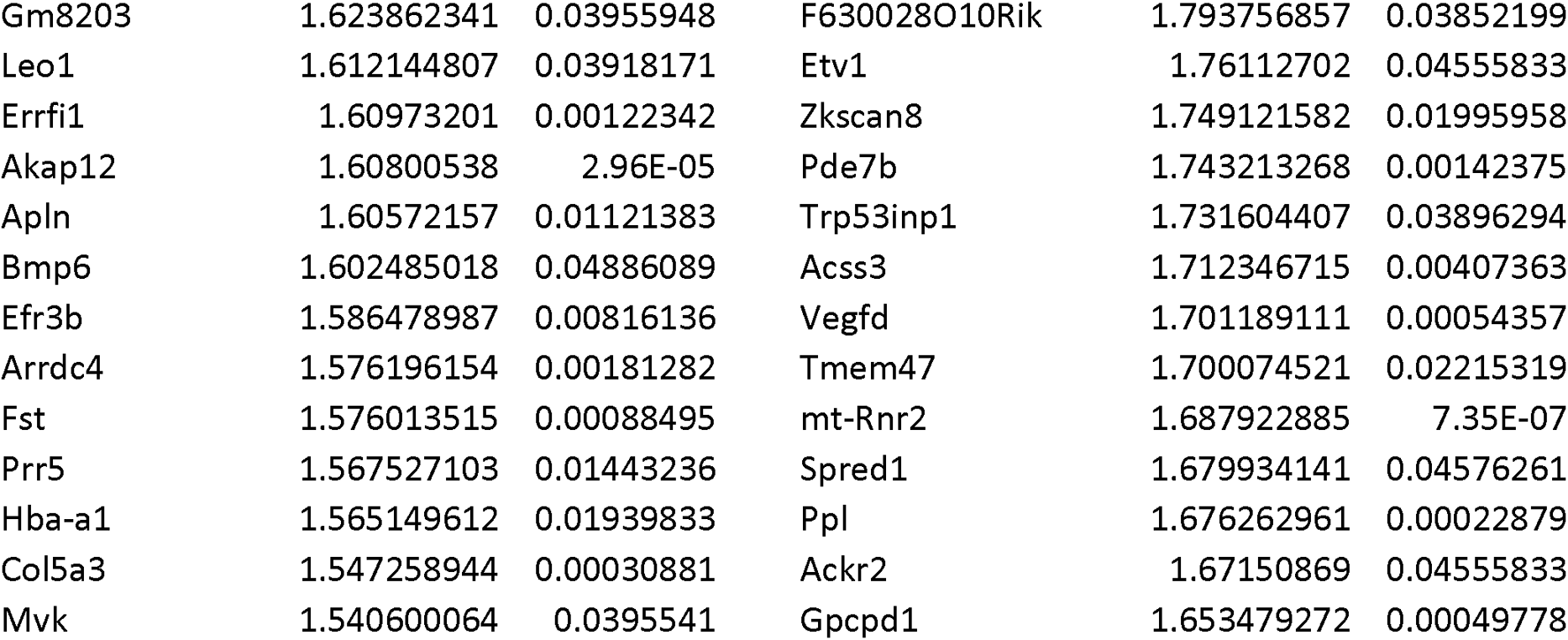
Top 50 Differentially expressed mouse genes between WIM and TIM.

**TABLE 2:**
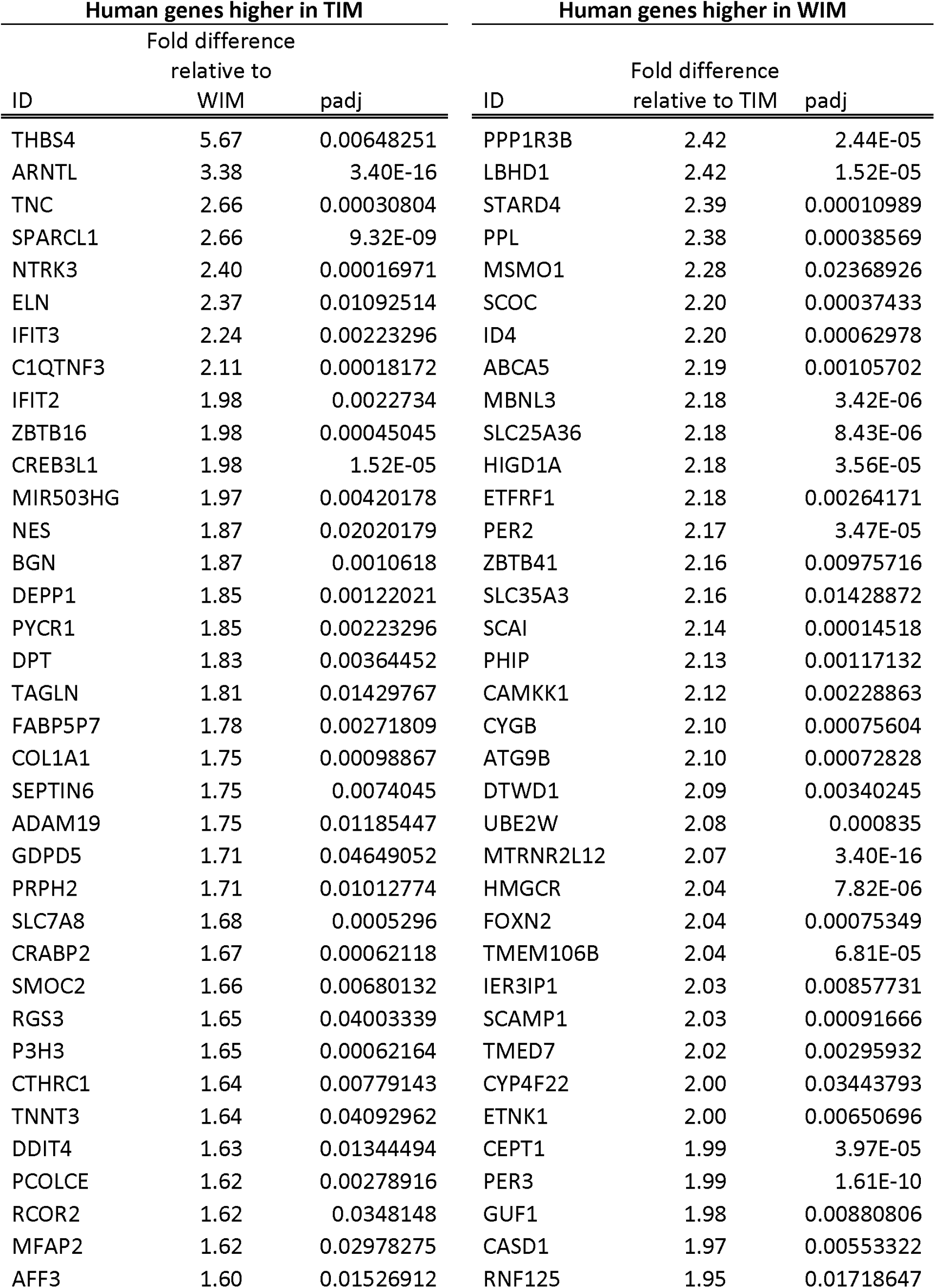
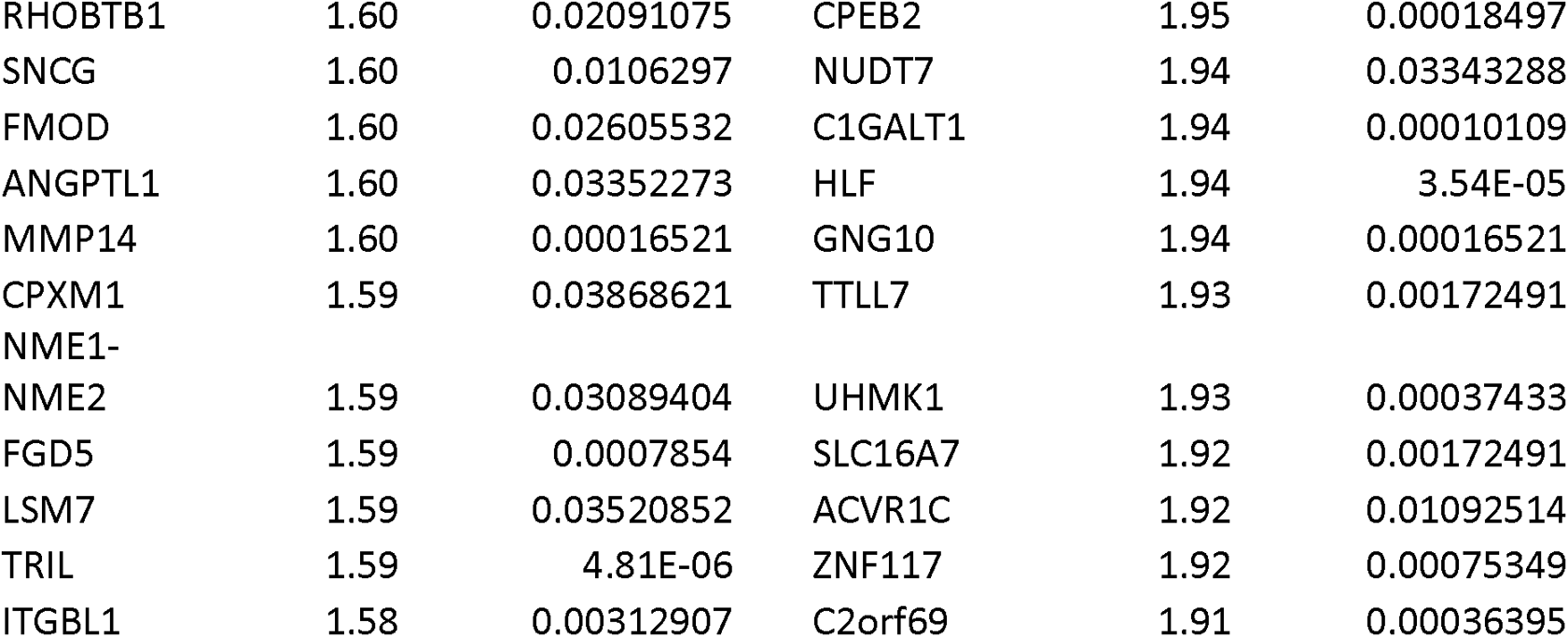
Differential expression of human genes between WIM and TIM.

**TABLE 3.** Top 50 genes differentially expressed between non-thermogenic and thermogenic adipocytes prior to implantation.

| Higher in thermogenic adipocytes prior to implantation |  |  | Higher in non-thermogenic adipocytes prior to implantation |  |  |
| --- | --- | --- | --- | --- | --- |
| Gene | foldChange | padj | Gene | foldChange | padj |
| C2CD4A | 155.34 | 0.002721933 | FABP7 | 183.54 | 6.7908E-10 |
| LINC00473 | 109.62 | 6.4997E-19 | LTF | 122.49 | 0.001425675 |
| PTPRN | 85.71 | 2.9956E-36 | KRT34 | 70.27 | 1.31375E-18 |
| FGF10 | 76.55 | 2.11699E-08 | KRT19 | 62.93 | 1.36279E-32 |
| CST2 | 68.88 | 5.98905E-06 | XIRP1 | 47.05 | 4.18785E-08 |
| AS3MT | 63.69 | 0.004661966 | PKD2L1 | 45.65 | 2.04601E-07 |
| GPR84 | 57.23 | 0.000211284 | TF | 43.82 | 3.99386E-05 |
| DLX5 | 46.37 | 4.77919E-07 | ANKRD1 | 32.84 | 2.29796E-19 |
| FZD10 | 45.52 | 0.003220422 | CLEC3B | 31.37 | 2.41762E-18 |
| ADCY4 | 43.43 | 1.80533E-35 | PKD1L2 | 31.28 | 5.99291E-18 |
| PTPN5 | 39.58 | 0.020773756 | MYEOV | 26.82 | 8.90154E-07 |
| FZD10-AS1 | 37.75 | 0.003260077 | KRT33B | 26.27 | 0.000146636 |
| HPD | 37.19 | 3.09808E-27 | PSG1 | 24.87 | 0.000828756 |
| FAM71E2 | 36.56 | 0.031407222 | AZGP1P1 | 23.52 | 5.06026E-05 |
| SYT6 | 35.78 | 0.002696626 | MYO18B | 23.44 | 0.000621079 |
| SEMA6B | 32.85 | 5.63876E-06 | C6 | 21.98 | 2.16083E-23 |
| VMO1 | 32.57 | 6.1947E-26 | DCSTAMP | 21.30 | 9.3728E-06 |
| CLEC2L | 32.41 | 0.000520436 | C8orf34 | 20.81 | 6.19938E-20 |
| PCSK2 | 30.73 | 1.94392E-05 | TRIML2 | 19.94 | 4.84051E-05 |
| SERTM1 | 30.71 | 1.12166E-05 | ITIH1 | 18.99 | 4.18994E-11 |
| ATP2B2 | 29.77 | 0.000200102 | SLC14A1 | 18.79 | 3.43563E-08 |
| SPINK6 | 29.41 | 1.73403E-12 | SERPINB2 | 18.44 | 1.18779E-18 |
| LGSN | 28.64 | 1.00209E-06 | LOC286189 | 18.12 | 0.007948722 |
| DLK1 | 28.63 | 2.96096E-16 | NLRP10 | 18.02 | 1.91549E-22 |
| CD86 | 28.45 | 0.002953203 | BIRC3 | 17.29 | 2.43275E-29 |
| CYSLTR2 | 27.31 | 0.006812095 | C1orf95 | 17.20 | 3.61531E-30 |
| LRP1B | 26.82 | 1.1216E-12 | HSPA7 | 16.88 | 7.28876E-06 |
| PCYT1B | 25.69 | 3.22807E-06 | NMUR1 | 16.85 | 1.23662E-22 |
| MTUS2 | 24.87 | 3.04361E-07 | KIAA0040 | 15.98 | 2.70823E-14 |
| DNASE1L3 | 24.69 | 2.69677E-13 | KY | 15.89 | 3.32077E-16 |
| VANGL2 | 24.51 | 2.30367E-10 | KRTAP2-3 | 15.51 | 1.51909E-20 |
| CACNG7 | 24.49 | 0.03961277 | NTSR2 | 15.48 | 7.55738E-05 |
| FNDC1 | 22.10 | 4.05767E-49 | S100B | 15.28 | 3.48826E-08 |
| KLRC2 | 22.03 | 0.002213239 | RELN | 15.06 | 1.97867E-14 |
| CCNJL | 22.00 | 0.000481462 | LOC101929371 | 14.11 | 4.47959E-28 |
| BAI3 | 19.88 | 0.002551949 | ACHE | 14.06 | 3.14035E-62 |
| TMEM190 | 19.72 | 0.000576069 | FOSB | 13.96 | 2.33018E-07 |
| MMP1 | 19.57 | 1.15956E-05 | TM4SF4 | 13.43 | 0.003571504 |
| SNAP25-AS1 | 17.46 | 1.17292E-05 | FCGR3A | 12.99 | 0.000292039 |
| HUNK | 17.38 | 3.21395E-11 | LOC100507065 | 12.72 | 7.94747E-22 |
| TNFSF11 | 17.06 | 2.48986E-05 | APOC1 | 12.72 | 4.57228E-16 |
| C3AR1 | 16.56 | 1.50178E-05 | HPR | 12.49 | 0.00064666 |
| ASB4 | 16.46 | 0.016524355 | FCRLA | 12.37 | 1.77321E-28 |
| MSI1 | 16.43 | 0.00036933 | CLCA2 | 12.21 | 2.82717E-15 |
| OGDHL | 15.97 | 3.55541E-11 | CD14 | 11.74 | 1.17228E-43 |
| ASPG | 15.45 | 3.26859E-06 | MARCO | 11.63 | 1.55074E-05 |
| KLRC3 | 14.78 | 0.000730806 | PFKFB1 | 11.60 | 2.8204E-21 |
| DYSF | 14.40 | 3.21559E-16 | LINC00161 | 11.49 | 0.012740702 |
| BTBD11 | 14.24 | 8.05332E-06 | PRKAG3 | 10.81 | 8.56138E-05 |
| CCDC177 | 14.00 | 0.00013086 | CCR1 | 10.66 | 5.71879E-19 |
| COL14A1 | 13.51 | 2.92944E-26 | AZGP1 | 10.62 | 8.36365E-10 |

**TABLE 4.**
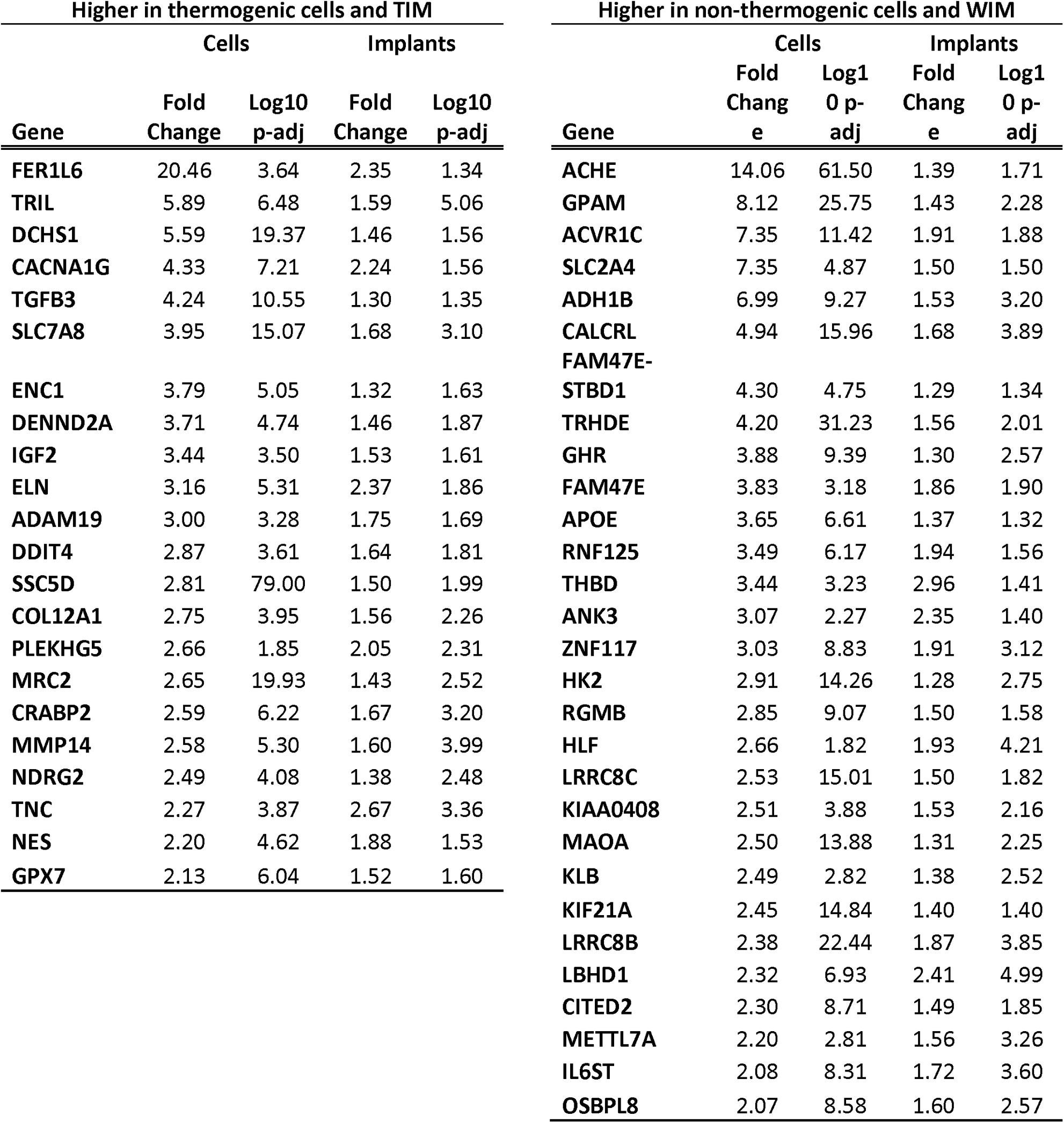
Genes differentially expressed between non-thermogenic and thermogenic adipocytes that are maintained differentially expressed in WIM and TIM.

